# Growth Hormone Accelerates Recovery From Acetaminophen-Induced Murine Liver Injury

**DOI:** 10.1101/2023.04.17.537197

**Authors:** Elissa Everton, Mercedes Del Rio-Moreno, Carlos Villacorta-Martin, Pushpinder Singh Bawa, Jonathan Lindstrom-Vautrin, Hiromi Muramatsu, Fatima Rizvi, Anna R. Smith, Ying Tam, Norbert Pardi, Rhonda Kineman, David J. Waxman, Valerie Gouon-Evans

## Abstract

**Background and Aims:** Acetaminophen (APAP) overdose is the leading cause of acute liver failure, with one available treatment, N-acetyl cysteine (NAC). Yet, NAC effectiveness diminishes about ten hours after APAP overdose, urging for therapeutic alternatives. This study addresses this need by deciphering a mechanism of sexual dimorphism in APAP-induced liver injury, and leveraging it to accelerate liver recovery via growth hormone (GH) treatment. GH secretory patterns, pulsatile in males and near-continuous in females, determine the sex bias in many liver metabolic functions. Here, we aim to establish GH as a novel therapy to treat APAP hepatotoxicity.

**Approach and Results:** Our results demonstrate sex-dependent APAP toxicity, with females showing reduced liver cell death and faster recovery than males. Single-cell RNA sequencing analyses reveal that female hepatocytes have significantly greater levels of GH receptor expression and GH pathway activation compared to males. In harnessing this female-specific advantage, we demonstrate that a single injection of recombinant human GH protein accelerates liver recovery, promotes survival in males following sub-lethal dose of APAP, and is superior to standard-of-care NAC. Alternatively, slow-release delivery of human GH via the safe nonintegrative lipid nanoparticle-encapsulated nucleoside-modified mRNA (mRNA-LNP), a technology validated by widely used COVID-19 vaccines, rescues males from APAP-induced death that otherwise occurred in control mRNA-LNP-treated mice.

**Conclusions:** Our study demonstrates a sexually dimorphic liver repair advantage in females following APAP overdose, leveraged by establishing GH as an alternative treatment, delivered either as recombinant protein or mRNA-LNP, to potentially prevent liver failure and liver transplant in APAP-overdosed patients.

## INTRODUCTION

Acetaminophen (acetyl-para-aminophenol, APAP) is consumed by over 60 million Americans weekly, making it the most frequently used analgesic and antipyretic in the US. Yet, APAP overdose is the most common cause of acute liver failure in the US^1^. Hepatotoxicity occurs when excess APAP overwhelms the urinary excretory pathway and is activated by the cytochrome P450 CYP2E1 metabolic pathway, generating the toxic metabolite N-acetyl-p-benzoquinone imine (NAPQI)^2–4^. Glutathione (GSH) binds and neutralizes NAPQI to a non-toxic metabolite; however, under overdose conditions, the rate of GSH synthesis is insufficient, leading to hepatocyte cell death^5–7^. The only treatment currently available is the administration of oral and intravenous Nacetyl cysteine (NAC), a precursor for GSH. However, NAC effectiveness rapidly diminishes within ten hours after APAP overdose^6–8^, when organ toxicity is frequently still asymptomatic^9^. Here, we address this unmet clinical need by taking advantage of the clinical^10–16^ and preclinical^17–20^ observation that females are more resistant to APAP toxicity, and in general to most liver diseases than males. Sex-specific hormones and their receptors drive sexual dimorphism in many liver processes in mice^18,21–24^ including hepatocyte proliferation, GSH synthesis, drug metabolism, and cell cycle inhibition. Specifically, gonadal steroid-controlled pituitary GH secretory patterns, pulsatile in males and persistent/near-continuous in females^25^, a pattern also observed in rodents^26–28^, is the basis of sex bias in liver function, including metabolic activity and disease susceptibility^29^. Namely, STAT5b has been reported to be an essential transcriptional regulator of the sex-biased actions of GH in the liver contributing to 90% of differences in liver transcriptome^30^. Consistent with prior literature, we demonstrate that females are more resistant to APAP toxicity than males. Single-cell RNA sequencing (scRNA-seq) analyses reveal that female hepatocytes express significantly higher levels of GH receptor (GHR) and GH pathway activation in comparison to male cells. As a result, we leverage this sexually dimorphic response to APAP and demonstrate the therapeutic benefit of recombinant GH protein treatment over the standard-of-care NAC treatment to significantly and rapidly repair the liver in both sexes following sex-specific sub-lethal doses of APAP, albeit in a milder fashion in females. Importantly, our study also introduces the use of nucleoside-modified mRNA encoding GH complexed to lipid nanoparticles (mRNA-LNP) for safe and slow-release delivery to the liver as an alternative to recombinant protein bolus for treatment of APAP overdose, a delivery platform recently validated in mRNA-based COVID-19 vaccines.

## METHODS

### Acetaminophen-induced liver injury mouse model

All mice used for the liver injury models are 10–12 week-old males and females inbred C57BL/6J from Jackson Laboratory. All animal studies were approved by the Boston University IACUC and were consistent with all local, state, and federal regulations as applicable. Mice were housed under standard conditions with a 12hour day/night cycle, in a pathogen-free environment with access to food and water ad libitum. Acetaminophen (APAP; Spectrum Chemical Manufacturing Corporation cat#AC100125GM) was dissolved in sterile PBS to 20 mg/ml concentration at 56°C, then cooled to room temperature. APAP was injected intraperitoneally at 10am for all experiments to remove circadian rhythm as a potential variable, following the Whitten effect^31–33^ in which female mice were placed in cages with soiled male bedding 48 hours prior to injury, and a 14-hour fast, to allow synchronization of estrus cycle which may influence APAP-induced liver injury and repair, and to normalize glutathione levels from the diet that may influence liver damage among the mice, respectively. Mice were maintained on normal chow diet and water ad libitum after APAP injections. Mice were euthanized as per IACUC regulation using isoflurane and cervical dislocation. Liver tissue and blood serum were collected from each mouse at the respective day of sacrifice.

## RESULTS

### APAP-induced liver injury and subsequent repair are sexually dimorphic

In dose response experiments in both sexes using a range of APAP doses from 300-650 mg/kg, previously validated by others mostly in male mice^34,35^, we found that 400 mg/kg APAP induces injury in both sexes without lethality up to 96 hours after APAP administration (Figure 1). Female mice were subjected to the Whitten effect, by replacing the female cage bedding with soiled male bedding to normalize estrus cycles of female mice^31–33^, as variability in estrogen levels may impact liver regeneration^10,12,36–39^. All mice were then fasted for 14 hours prior to APAP injection to bring liver metabolism to a baseline level and thus help normalize injury^34,35^ (Figure 1A). Sexually mature male and female mice were injected intraperitoneally (IP) with APAP or PBS vehicle control, and euthanized 12, 24, 48, and 96 hours later. Serum alanine aminotransferase (ALT) levels, indicative of liver damage, were consistently higher in males than in females at all time points post-24 hours (Figure 1B). Acute central vein area necrosis was seen in both sexes 24 hours after APAP injection, consistent with the localization of CYP enzymes metabolizing APAP^3,40^. Necrosis was more extensive in male livers and progressively increased over time, whereas female livers fully recovered by 48 hours (Figure 1C). Cell death, as indicated by TUNEL assay, showed a similar sex difference (Figure 1D). Consistent with the histological data, serum bilirubin persisted for 96 hours in males but decreased after 48 hours in females, supporting recovery in females that is not seen in males (Figure 1E). Thus, males are more susceptible than females to APAP hepatotoxicity as characterized by progressive, unresolved necrosis and apoptosis that most likely prevent liver regeneration.

**Figure 1:**
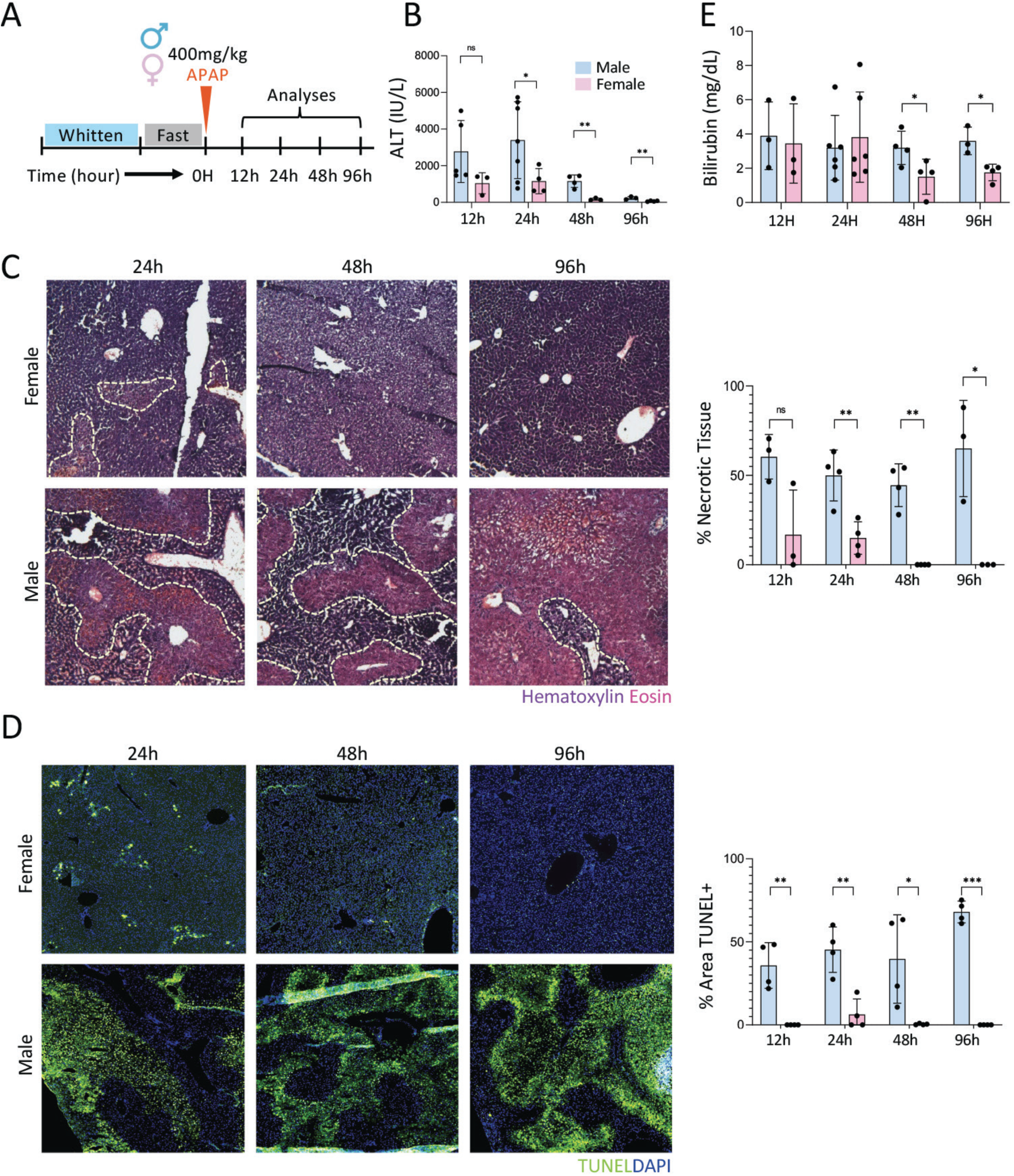
Liver necrosis and cell death are sexually dimorphic following administration of equivalent doses of APAP. A: Following the Whitten effect and fasting to normalize estrogen and glutathione levels, respectively, male and female 12-week-old C57BL/6J mice were injected with 400 mg/kg APAP intraperitoneally and followed for 96 hours using the injury scheme outlined. B: Serum alanine aminotransferase (ALT) levels at each sacrifice time post-APAP. C: Hematoxylin & eosin staining of representative male and female livers at each time point, shown at 40X magnification. Bar graph on the right quantifies tissue area covered by necrosis (H&E). D: TUNEL staining of representative male and female livers at each time point, quantifying both cell apoptosis and necrosis, shown at 40X. Bar graph on the right quantifies tissue area covered by cell death (TUNEL+). E: Serum bilirubin in each sex at each time point. N=3-6 mice per sex/time point, with 2-3 lobes averaged for histological quantification – one male mouse died before 96 hours. Statistics calculated via two-sided student’s t test for unpaired comparisons; p-values *<0.05, **<0.001, ***<0.0001.

### Growth hormone receptor pathway activation is more enriched in female livers

To identify druggable sexually dimorphic pathways activated in females that promote liver repair after APAP overdose, we performed a scRNA-seq analysis for male and female liver cells, treated with PBS control or 400 mg/kg APAP at 48 hours post-injection (Figure 2A). Single liver cells were collected as per a modified protocol^41^, to obtain a mix of ∼40% hepatocytes (Hep) and ∼60% nonparenchymal cells (NPC) including immune cells, endothelial cells (EC), and fibroblasts/hepatic stellate cells (HSC). Following scRNA-seq of the four samples, data were combined (Figure 2B) or grouped based on treatment (Figure 2C), analyzed using the Seurat package, and imported into SPRING software^42^ (Figure 2B,C). Interestingly, hepatocytes (Heps) and endothelial cells (ECs) segregated separately based on their sex prior to injury, then transcriptionally shifted closer to the other sex following injury as indicated with the SPRING plots (Figure 2B, C) and ENRICHRbased pathway analyses (Supplemental Figure 1A). Bulk RNA-seq analyses have reported sexual dimorphism of liver transcriptomes^43–45^, and both periportal and pericentral hepatocytes were recently identified as the main cell types contributing to the disparity^46^. Our single-cell data reveal that not only hepatocytes harbor distinct transcriptomes based on sex, but ECs do as well. Given the known sex differences in GH secretion patterns and their impact on liver metabolism^29,43,47,48^, we specifically examined the GHR pathway in single cells. Violin plots show that female Heps express significantly higher levels of GHR than male cells, both before and after APAP injection. Only a small percentage of ECs express GHR in both sexes (Figure 2D). Importantly, the enrichment in BioCarta gene sets related to GH/GHR pathway activation was globally greater in Heps and ECs from female mice than from male mice after APAP injection (Figure 2E), yet levels of a set of specific GH/GHR-pathway activation related genes was also significantly greater in females cells compared to their male counterparts prior to APAP injury (Supplemental Figure 1B includes genes from the BioCarta GH pathway activation; Supplemental Figure 1C includes additional genes key in the GH pathway activation). These data illustrate the association between GH/GHR pathway activation and accelerated liver regeneration seen in females, suggesting that given the known GH-dependent sexual dimorphism of liver metabolism, administration of GH could improve recovery after acute liver injury caused by APAP overdose.

**Figure 2:**
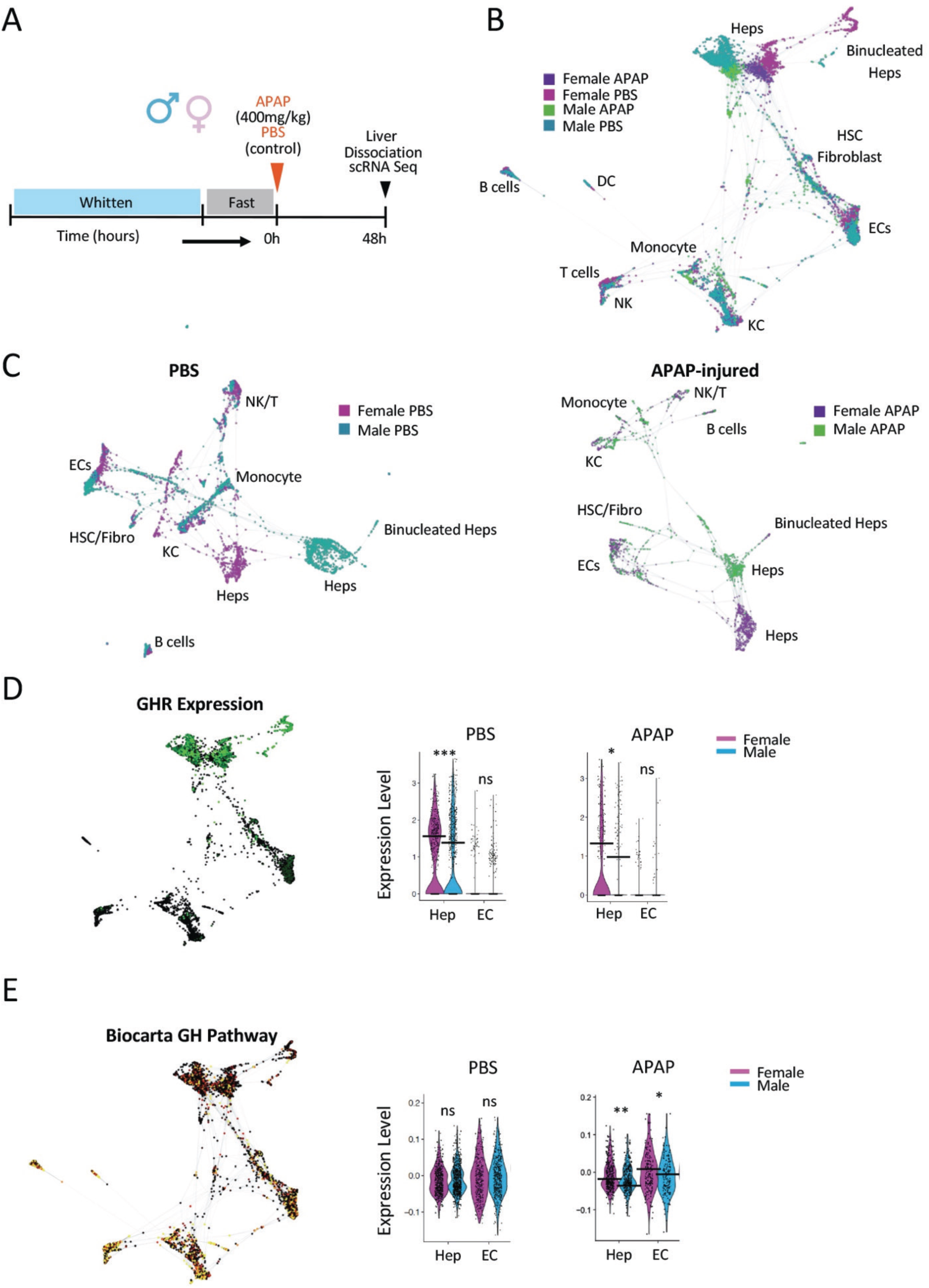
Male and female livers exhibit distinct transcriptional responses to acetaminophen-induced liver injury. A: Male and female mice were given equivalent doses of 400 mg/kg APAP or PBS vehicle control post-fast and Whitten (for females), then livers were perfused in situ and dissociated at 48 hours post-APAP treatment for single-cell RNA sequencing analysis. B: All 4 datasets combined on a single SPRING plot (4,821 cells), with 8 different cell type clusters resolved. Hepatocytes and endothelial cells (ECs) exhibit sexually dimorphic clustering prior to APAP injury. C: Subclusters of combined dataset, separated by treatment, with female cells in pink and purple, and male cells in teal and green. D: Growth hormone receptor (GHR) expression of combined SPRING plot, and violin plots of PBS-treated and APAP-treated hepatocytes (Hep), endothelial cells (EC) showing differential expression of GHR between males (blue) and females (pink). E: Growth hormone (GH) pathway enrichment of Biocarta gene set in SPRING and violin plots of PBS-treated and APAP-treated Heps and ECs showing differential enrichment of GH pathway between males (blue) and females (pink). DEG of 0.25 used for differential expression resolution. P-values *<0.05, **<0.001, ***<0.0001.

### A single injection of recombinant human GH accelerates liver repair after APAP overdose

As a first approach to investigate the clinical benefit of GH, we tested the efficiency of a single injection of human recombinant GH to promote liver repair in both sexes after female and male livers were similarly injured with APAP. To achieve this, male and female mice were injected with sex-specific sub-lethal doses of APAP selected to achieve a similar level of toxicity in each sex, namely 400 mg/kg for males and 600 mg/kg for females. Eight hours later mice were given a single subcutaneous injection of recombinant human GH (2.5 mg/kg) or control PBS, then analyzed 24 and 48 hours after APAP overdose, and observed for up to 6 days to compare survival between the sexes and treatments (Figure 3A). We confirmed that the sex-specific APAP doses chosen induced a similar degree of liver injury in both sexes 12 hours after APAP overdose, when the injury is the greatest (Supplementary Figure 2A), as shown with no significant difference in serum ALT levels and necrotic areas, although the expansion of apoptotic TUNEL+ area was significantly lower in females. To determine the minimum effective dose of GH, we tested doses (0.5, 2.5, 5, 10 mg/kg) ranging from 0.5 mg/kg^49,50^ to 10 mg/kg^51,52^ (Supplementary Figure 2B), including the average dose of 1 mg/kg, used for adolescents treated daily with GH for short stature therapy^53^ (which in mice translates to ∼12.3 mg/kg using an inter-species dose conversion factor^54^). We found that 2.5 mg/kg GH was the minimum dose that significantly decreased necrosis in both sexes and serum ALT levels in males as compared to PBS controls (Supplementary Figure 2C,D); this dose was therefore used in all subsequent studies. The highest GH dose, 10 mg/kg, was found toxic in both sexes as indicated by high serum ALT levels in males and expansion of necrotic tissue in both sexes similar to that found in PBS controls (Supplemental Figure 2C–E). 2.5 mg/kg of GH significantly lowered serum ALT levels 48 hours after APAP administration in males compared to PBS-treated males (Figure 3B), an effect that was not seen in females, most likely due to ALT levels already being lowered in females at both time points. The beneficial effect of GH in males was further illustrated by the sharp and significant GH-dependent decreases in necrotic areas at both time points and in apoptotic TUNEL+ area 24 hours post-APAP (Figure 3C, D; Supplemental Figure 3A, B). The effect of GH in females was less pronounced, most likely due to the continuous endogenous GH secretion already present in females, yet injection of GH significantly and specifically decreased the necrotic areas 24 hours post-APAP compared to PBS-treated females (Figure 3C, Supplemental Figure 3A). Six-day survival data showed that of 5 APAP-treated mice in each sex, 1 PBS-treated female died (80% survival) and 2 PBS-treated males died (60% survival), while GH increased mouse survival up to 100% for females and 80% to males. Overall, GH treatment sharply and significantly decreases liver injury and accelerates regeneration when given 8 hours post-APAP in both sexes, with a more substantial effect in males.

**Figure 3:**
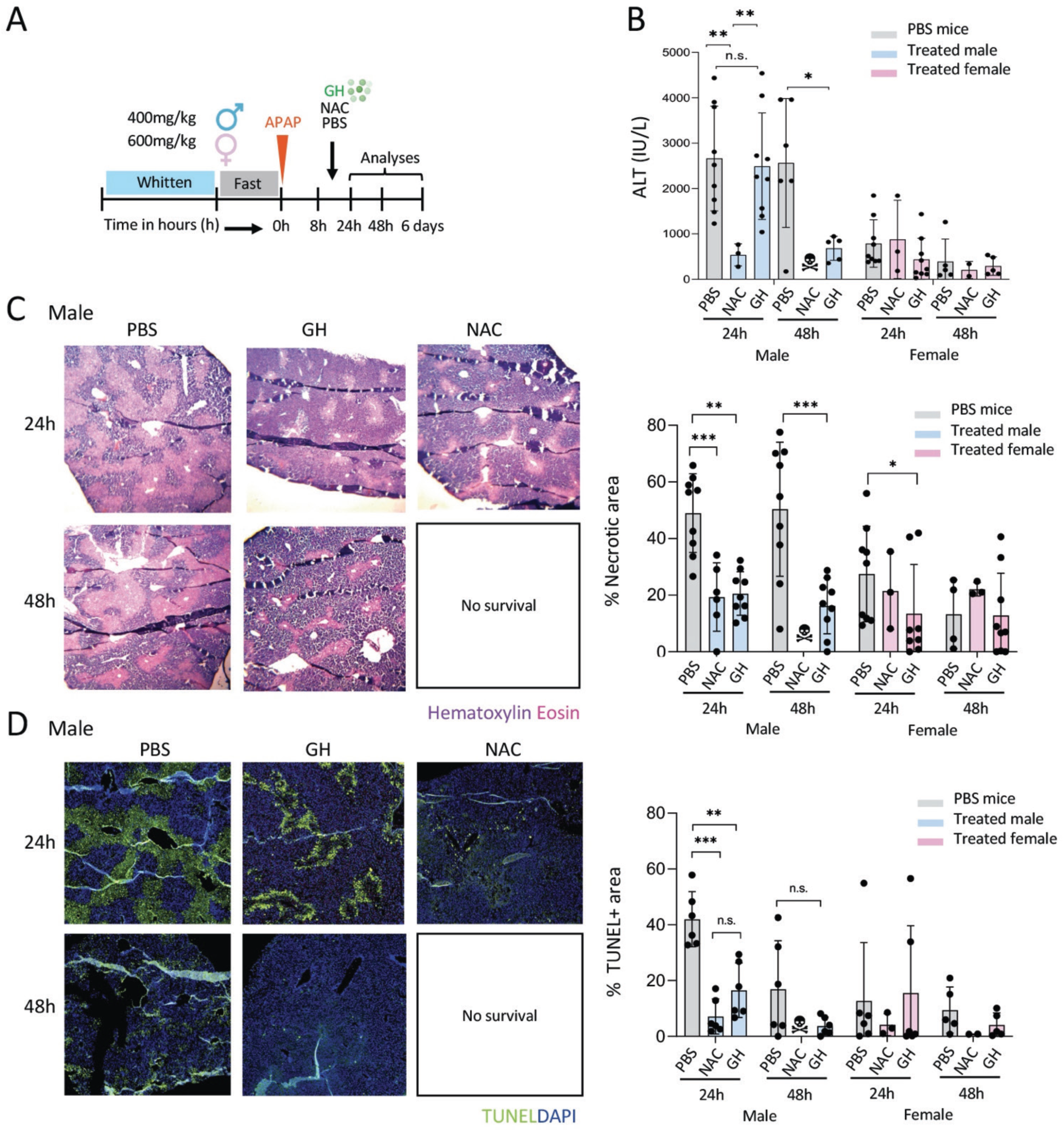
Exogenous human growth hormone treatment promotes liver recovery from APAP-induced injury and more efficiently than NAC. A: Injury and treatment scheme consisting of fast and Whitten effect (females), then differential severe doses of APAP (400 mg/kg for males, 600 mg/kg for females), followed by treatment of 2.5 mg/kg GH, 1000 mg/kg NAC, or equivalent volume of PBS vehicle control 8 hours after APAP injection. N=5-9 mice per sex/treatment/time point (GH and PBS) or N=3 per sex/time point (NAC), with 2-3 lobes averaged per mouse for histological quantification. B: Serum ALT levels of GH-treated, NAC-treated, and PBS-treated male and female mice 24 and 48 hours after severe dose of APAP. C: Representative H&E stains of male mouse livers per treatment, and time point post-APAP at 40X magnification, and graph quantifying % necrotic tissue in males and females. D: Representative TUNEL stains of male mouse livers per treatment, and time point post-APAP at 40X magnification and graph quantifying % TUNEL+ tissue in males and females. Statistics calculated via two-sided student’s t test for unpaired comparisons; p-values *<0.05, **<0.001, ***<0.0001.

### Human growth hormone treatment accelerates liver recovery better than administration of NAC

For potential clinical translation, we compared the efficacy of GH treatment to that of the clinical standard-of-care treatment with NAC to accelerate liver regeneration post-APAP overdose in the same sets of experiments (Figure 3A–D; Supplemental Figure 3). Sex-specific sub-lethal doses of APAP were administered to mice, which were then treated 8 hours later with either GH (2.5 mg/kg), NAC (1000 mg/kg, the average dose reported in murine APAP overdose to mitigate liver injury in murine APAP overdose studies^55–59^), or PBS control. Impressively, all NAC-treated males died by 48 hours post-APAP, while all males treated with GH or control PBS survived (Supplemental Figure 3C) suggesting a toxic effect of NAC in males. In contrast, all females survived regardless of the treatment. For the male mice that survived 24 hours post-APAP, NAC significantly diminished necrotic and TUNEL+ areas, although these mice did not survive by 48 hours post-APAP (Figure 3C, D). This indicates that the beneficial effect of NAC is short-lived when compared to that of GH, which significantly decreased necrotic and apoptotic areas in males and necrotic areas in females compared to PBS (Figure 3C, D; Supplemental Figure 3A). The deleterious effect of NAC treatment may be due to an anaphylactoid reaction, which has been reported in up to 18% of patients receiving intravenous NAC^60^, or to an increase in clotting time^61^. In females, NAC did not show the beneficial effect of reducing necrotic tissue seen with GH (Figure 3C, D, Supplemental Figure 3A, B). Altogether, when given 8 hours post-APAP, GH is more efficient in accelerating liver repair in both sexes, as compared to a short-lived beneficial effect of the standard-of-care NAC seen only in males. This suggests that GH may have complementary clinical benefit for APAP-overdosed patients that present late in the ER after NAC loses its efficacy.

### Constitutive activation of the downstream GH/GHR pathway mediator STAT5b rescues males from APAP hepatotoxicity

To further validate the clinical benefit of GH treatment in accelerating liver repair, we investigated whether constitutive activation in hepatocytes of the known downstream GH/GHR pathway mediator, transcriptional regulator STAT5b^30,47,48^ (STAT5bCA) responsible for the major effects of GH on sex differences in the liver^48,62^, would have a protective role prior to APAP injury. Male mice, in which GH treatment was the most significant, were injected with AAV8-TBG-STAT5bCA to induce TBG promoter-driven hepatocyte-specific expression^63–65^ of the mutated STAT5b (STAT5bCA) which is constitutively active even in the absence of GHR activation with GH^30^ (Figure 4). STAT5bCA has been reported to mimic the response of endogenous liver STAT5b to the female, persistent circulating GH profile and thereby feminizes gene expression in the liver^30^. Seven days later, mice were fasted, given 400 mg/kg APAP, and sacrificed 48 hours post-APAP treatment (Figure 4A). Expression and function of STAT5bCA in hepatocytes through AAV8-TBGTAT5bCA injection was validated by a significant increase in plasma IGF1 concentration as previously reported^30^ compared to control AAV8-Null-treated mice 6 days post-AAV8 injection (Figure 4B). Remarkably, while only one of the 6 AAV8-Null-treated control mice survived after 48 hours, all 6 AAV8-STAT5bCA-treated mice survived (Figure 4C), although necrosis and apoptosis were still visible (Figure 4D). These findings demonstrate that persistent activation of hepatocytic STAT5b, as normally occurs in female liver, confers a striking protective effect from APAP-induced liver injury, and further support the clinical benefit of GH administration to rapidly promote liver recovery from APAP hepatotoxicity.

**Figure 4:**
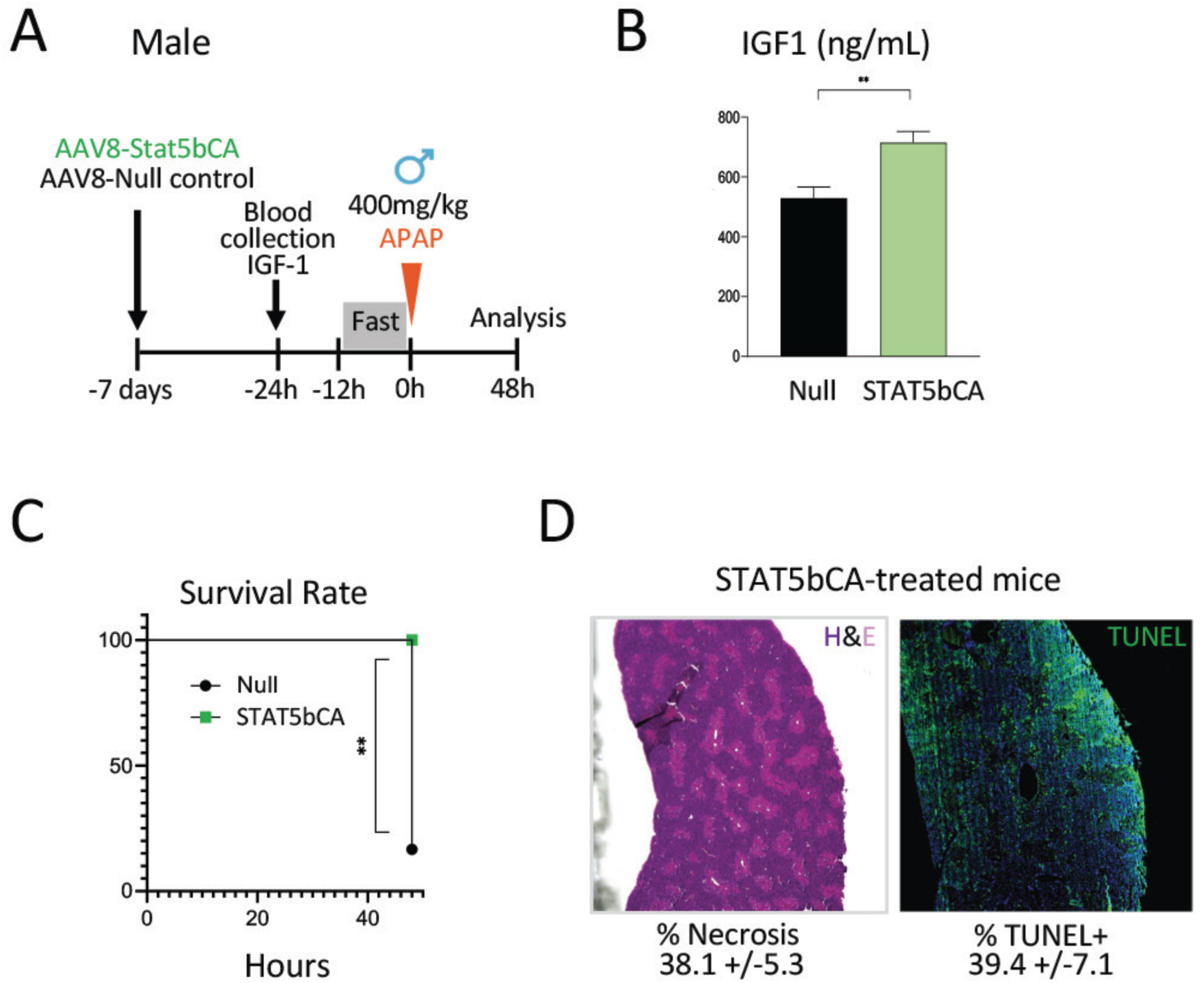
Constitutive activation of the downstream GH/GHR pathway mediator STAT5b rescues males from APAP hepatotoxicity. A: Pre-treatment scheme with AAV8-TBGSTAT5bCA given 7 days prior to 400 mg/kg severe APAP injury, in C57BL/6J male mice, and sacrificed at 48 hours post-APAP administration. B: Plasma murine IGF1 concentration measured by ELISA 6 days post-AAV8 injection in Null(black) and STAT5bCA-injected (green) mice. C: Survival rate for STAT5bCA-(green square) and Null-injected mice (black x). D: Representative H&E stain and TUNEL stain of STAT5bCA-injected mouse livers, with average of % necrotic tissue and % TUNEL+ cell area and standard error for treatment group. N=6 mice/treatment. Statistics calculated via two-sided student’s t test for unpaired comparisons; p-values *<0.05, **<0.001, ***<0.0001.

### Nucleoside-modified mRNA-LNP delivery of GH induces sustained GH expression over the course of liver injury and promotes recovery from APAP overdose

We next investigated the efficacy of nucleoside-modified mRNA encoding human GH encapsulated in lipid nanoparticles (GH mRNA-LNP) for slow-release delivery to the liver as an alternative to recombinant GH protein bolus. mRNA-LNP is a liver-targeted delivery technology that we recently implemented to express regenerative factors in the liver to promote liver regeneration and treat features of murine acute and chronic liver disease^66^. mRNA-LNP is a safe, non-integrative and non-immunogenic technology that has been widely validated in the form of current mRNA-LNP-based COVID-19 vaccines, and allows for robust yet transient expression of proteins with slow-release in the liver over the course of the injury induced by APAP. In this experiment, mice were injured with sex-specific sub-lethal doses of APAP, treated with either 10 μg of GH mRNA-LNPs or with control luciferase (Luc) mRNA-LNPs injected retro-orbitally 8 hours after APAP overdose, and then analyzed 24 and 48 hours after the overdose (Figure 5A). We first verified the efficient expression of human GH in mouse livers 5 hours post-injection (Figure 5B). A large increase in serum human GH was also seen, as the transfected hepatocytes ultimately secrete hGH in the blood stream as shown previously with other secreted factors delivered with mRNA-LNP^66^. Over time, serum human GH levels decrease, but are still detectable during the first 2 days post-injection, which represents the critical time during which APAP-induced injury occurs, and thus when GH is the most needed (Figure 5B). Remarkably, a single dose of GH mRNA-LNP rescued males from death, which occurred by 48 hours post-APAP in all control Luc mRNA-LNP-treated mice (Figure 5C), which was also associated with lower levels of ALT (Figure 5D). Though, 48 hours was likely too early to observe improvement of the liver tissue architecture (Figure 5E, F). In females, GH mRNA-LNP treatment showed no significant benefit, as recovery occurred spontaneously by 48 hours. This was not surprising, as serum human GH levels found in mice 5 hours post mRNA-LNP injection were 30-fold lower than serum levels extrapolated from a single injection of 2.5 mg/kg of recombinant GH, as tested in Figure 3. This suggests that increasing doses of slow-release GH mRNA-LNP may not only rescue male mice from death, but may further accelerate recovery in both sexes, as an alternative delivery platform to a bolus injection of recombinant GH protein.

**Figure 5:**
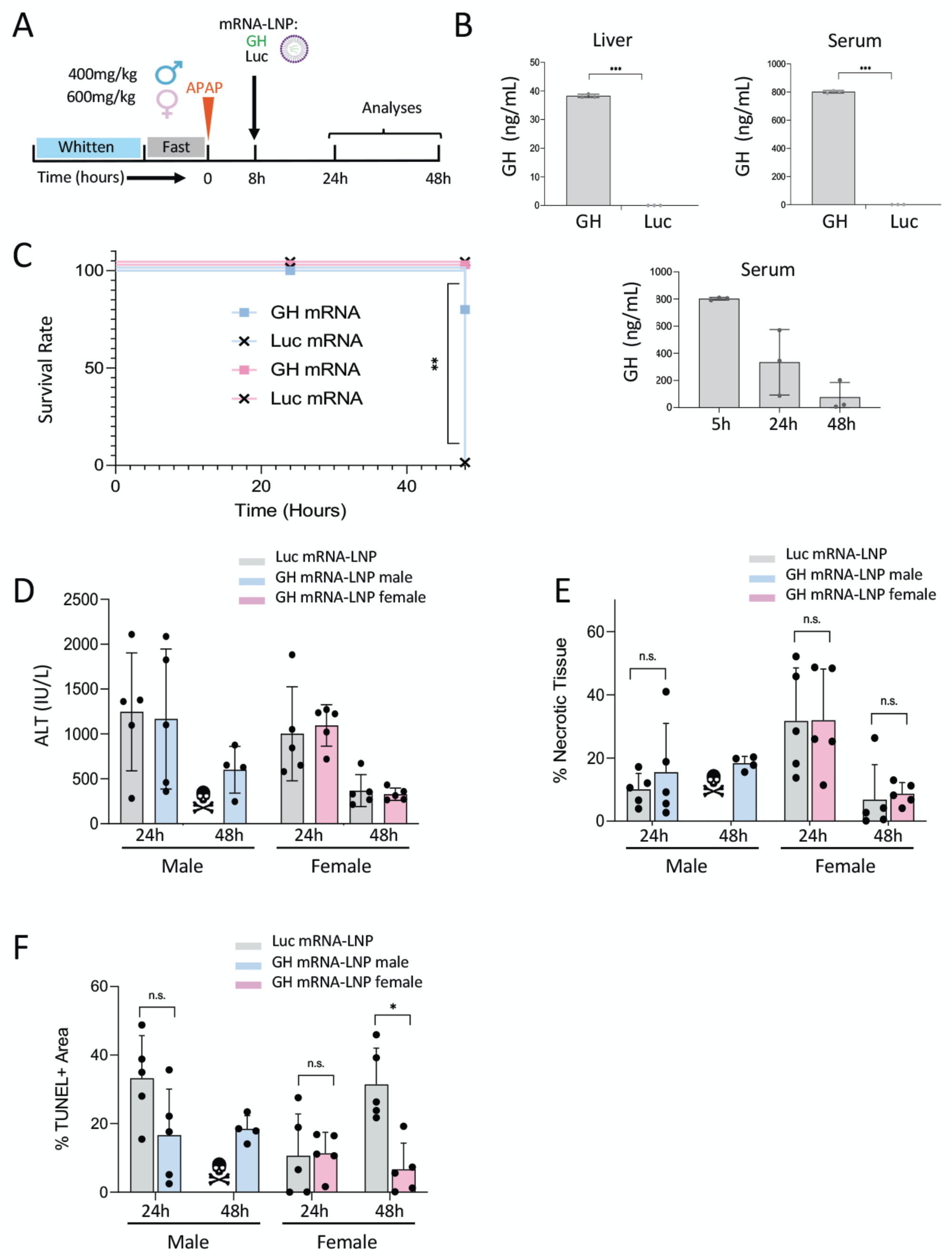
GH mRNA-LNP targets liver to promote recovery throughout the length of the APAP-induced acute injury. A: C57BL/6J mice were given sub-lethal doses of APAP (400 mg/kg for males, 600 mg/kg for females), and 8 hours later were injected retro-orbitally with 10 μg of human GH mRNA-LNP or negative control Luc mRNA-LNP, then analyzed 24 and 48 hours post-APAP. B: Upper panels: human GH protein levels in liver tissue homogenate (left) and serum (right) were measured 5 hours post-injection by ELISA. Lower panel: human GH protein levels in serum of mice were measured 5, 24 and 48 hours post GH mRNA-LNP injection by ELISA. N=3 mice per treatment/time point. C: Survival rate of GH mRNA-LNP-treated mice (squares; males in blue, females in pink) and Luc mRNA-LNP-treated mice (black x’s) 24 and 48 hours post APAP overdose. N=5 mice per sex/treatment/time point. Statistical analysis carried out using the Logrank Mantel-Cox test. D: Serum ALT of GH mRNA-LNP-treated females (pink) and males (blue) and Luc mRNA-LNP-treated (grey) mice 24 and 48 hours post APAP treatment. Note all male mice treated with Luc mRNA-LNP died by 48 hours post treatment (skull symbol). E: % necrotic liver tissue quantified from H&E stains in GH mRNA-LNP-treated (colored bars) and Luc mRNA-LNP-treated (grey bars) mice. F: % TUNEL+ area from TUNEL stain in GH mRNA-LNP-treated (colored bars) and control Luc mRNA-LNP-treated (grey bars) mice. Statistics for bar graphs calculated via two-sided student’s t test for unpaired comparisons; p-values *<0.05, **<0.001, ***<0.0001.

## DISCUSSION

NAC is the only current treatment for APAP overdose other than liver transplantation, and loses effectiveness ∼10 hours after APAP ingestion, when the symptoms of acute liver failure are frequently not yet evident^8^. Our study introduces a potential alternative therapeutic strategy to overcome this clinical unmet need for the many overdose cases that present late to the emergency room. This study reveals the sexually dimorphic response to APAP overdose and identifies GH/GHR/STAT5b as a sexually differentially activated and druggable pathway, that may ultimately be leveraged for the establishment of a GH-based therapeutic to accelerate recovery from APAP-induced liver injury. We demonstrate the efficiency of this therapy in mice using a single dose of GH, delivered as a recombinant protein or via mRNA-LNP injection and its superiority to NAC to mitigate injury, accelerate recovery, and promote survival.

The therapeutic capacity of GH to reverse APAP-induced hepatoxicity reported here is supported by earlier work demonstrating the ability of GH to stimulate liver cell proliferation and liver recovery via EGFR signaling in injured liver mouse models affected with partial hepatectomy or liver steatosis^67–70^. Consistent with the protective role of GH in APAP-induced hepatotoxicity, previous studies have shown that inhibition of GH-releasing hormone (GHRH) prior to APAP overdose increases APAP-induced mouse liver toxicity, while GHRH super-agonist partially reverses APAP toxicity^71^. Here, the pretreatment of male mice prior to APAP overdose with a constitutively active mutant form of the GH-activated transcription factor STAT5b, STAT5bCA, largely recapitulated the relative resistance to APAP seen in female, as compared to male mice and conferred striking protection from APAP overdose hepatotoxicity. Further study is required to determine whether the protective effect of STAT5bCA results from accelerated recovery from APAP hepatotoxicity or is due to a decrease in the extent of APAP-induced injury downstream of the overall feminization of the liver that results from AAV8-STAT5bCA treatment^30^.

GH is most likely not the only sex-dependent factor involved in the sexual dimorphism of APAP-induced injury and recovery, which was illustrated by the consistently slower liver recovery of GH-treated males as compared to females. Estrogens have been reported to have a direct impact on the GH/GHR pathway activation by reducing hepatic production of the GH-effector IGF-1 in response to GH^39^ and its sex-dependent pituitary secretory pattern. Interestingly, IGF-1 can also bind in a compensatory manner to estrogen receptor (ERα) under conditions of low systemic estrogen concentrations^72^. In addition to influencing the GH pathway, estrogen also contributes to liver function independently, for example, by increasing glutathione synthesis^21^ and by repressing mitochondrial expression of SH3 domain–binding protein that preferentially associates with Btk (SAB) in hepatocytes, thereby preventing liver injury from APAP overdose^18^. These protective effects of estrogen may contribute to the persistent sex differences in APAP-induced liver injury we observe even after GH administration, either directly via ERα expressed in hepatocytes, or indirectly, via its effect on pituitary GH secretion patterns^22–24^.

This study introduces GH delivery via intravenous injection of nucleoside-modified mRNA-LNP to induce production of GH protein throughout the duration of APAP-induced injury in order to accelerate recovery and increase survival. The use of mRNA-LNP to harness tissue regeneration following acute injury, such as that induced by APAP overdose, departs from its original applications for immunization in vaccines^73^ or protein replacement for diseases in which proteins are deficient or defective^74^. Given the short half-life of circulating GH after subcutaneous injection which ranges from 2 to 3 hours with a biological half-life of about 12 hours considering the downstream effects^75^, mRNA-LNP delivery of GH offers a wider timeframe of expression of at least 48 hours after a single injection, which should cover the duration of the acute liver damage caused by APAP. Following a single injection of GH mRNA-LNP (10 μg/mouse), the serum level of GH reached about 800 ng/mL, which is about 30x lower than was extrapolated when we injected recombinant GH protein. This suggests that the initial dose of GH mRNA-LNP we used in this study can be increased and carefully fine-tuned based on severity and persistence of liver disease. Therefore, delivery of GH via mRNA-LNP may be a promising alternative method to recombinant GH protein therapy for future clinical studies to treat acute liver failure.

In conclusion, this study demonstrates a sexually dimorphic liver repair process following APAP overdose benefitting female mice, which was leveraged by establishing GH as an alternative treatment to improve repair in males and accelerate it in females. This innovative advancement may prove critical in clinical translation for preventing liver failure and liver transplant for APAP-overdosed patients that present late in the ER, for whom NAC loses its therapeutic efficiency. In a clinical setting, GH administration could potentially be paired with standard-of-care NAC treatment as a complementary therapy activating broader mechanisms of repair to prevent liver failure or the need for transplant. Additionally, this study introduces the use of the safe non-integrative nucleoside-modified mRNA-LNP platform for delivering GH in a robust and sustained, yet controllable, transient manner, which was validated as safe with the recent mRNA-based COVID-19 vaccines^73,76–78^.

## ACKNOWLEDGEMENTS

This research was supported in part by the CTSI TL1TR001410 training grant in regenerative medicine (to EE and ARS) and the NIDDK F31DK127606-01A1 fellowship award (to EE), and by NIH grant DK121998 (to DJW). We would like to thank Amman Bhatti (Boston University) for her experimental contributions. We are grateful to Drs. Greg Miller and Marianne James of the CReM, supported by grants R24HL123828 and U01TR001810.

## AUTHOR CONTRIBUTIONS

Concept: Elissa Everton, Valerie Gouon-Evans, David Waxman, and Rhonda Kineman. Analysis: Elissa Everton and Mercedes del Rio-Moreno. Data acquisition: Elissa Everton and Mercedes del-Rio Moreno. Development of experimental materials: Hiromi Muramatsu, Norbert Pardi and Ying Tam. Bioinformatics: Pushpinder Singh Bawa, Carlos Villacorta-Martin, Jonathan Lindstrom-Vautrin, and Elissa Everton. Writing: Elissa Everton and Valerie Gouon-Evans. Editing: all authors.

## SUPPLEMENTAL INFORMATION

### Mouse treatments

Eight hours following APAP injury, mice were treated with recombinant human growth hormone (Invitrogen cat# PIRP10928) or NAC (Alfa Aesar cat# A1540914). GH and NAC were both diluted in sterile PBS. GH was diluted to a concentration of 1 mg/mL and given at dosage indicated, subcutaneously with a 0.5mL insulin syringe. NAC was diluted to a concentration of 50 mg/mL and given at 1000 mg/kg dosage, intraperitoneally with a 1mL insulin syringe.

For STAT5b activation experiments, 7 days prior to APAP injury, 10-12 week old male mice were treated with a single retroorbital injection of 7.5 × 10^10^ genome copies (GC) of AAV8-TBG-STAT5bCA^30^ or AAV8-TBG-Null diluted in 100μL of sterile PBS. To validate the activity of STAT5bCA, IGF1 was measured in plasma obtained from a lateral tail vein blood sample, collected the day prior to APAP injection.

For mRNA-LNP experiments, mRNA-LNPs were thawed and freshly diluted on ice in sterile Dulbecco’s Phosphate Buffered Saline (PBS) prior to each experiment. Mice were administered once 10μg mRNA-LNP in 50μl sterile PBS intravenously by retro-orbital injection under isoflurane anesthesia using a 1/2cc lo-dose insulin syringes (EXELINT) 8 hours after APAP injection.

### mRNA-LNP production

mRNA production was performed as described^79^. Briefly, sequences of the firefly luciferase and human growth hormone were codon-optimized, synthesized (GenScript), and cloned into an mRNA production plasmid. mRNAs were produced from linearized plasmids to contain 101 nucleotide-long poly(A) tails. m1Ψ-5’triphosphate instead of UTP was used to generate modified nucleoside-containing mRNA. Capping of the in vitro transcribed mRNAs was performed co-ranscriptionally using the trinucleotide cap1 analog, CleanCap. mRNA was purified by cellulose purification, as described^80^. mRNAs were analyzed by agarose gel electrophoresis and were stored frozen at –20°C. Cellulose-purified m1Ψ-containing RNAs were encapsulated in LNPs using a self-assembly process as previously described wherein an ethanolic lipid mixture of ionizable cationic lipid, phosphatidylcholine, cholesterol and polyethylene glycol-lipid was rapidly mixed with an aqueous solution containing mRNA at acidic pH^81^. The LNP formulation used in this study contains an ionizable cationic lipid (pKa in the range of 6.0–6.5, proprietary to Acuitas Therapeutics) /phosphatidylcholine / cholesterol / PEG-lipid^81,82^. The proprietary lipid and LNP composition are described in US patent US10,221,127 entitled “Lipids and lipid nanoparticle formulations for delivery of nucleic acids” (https://www.lens.org/lens/patent/183-348-727-217-109)6. They had a diameter of ∼80 nm as measured by dynamic light scattering using a Zetasizer Nano ZS (Malvern Instruments Ltd, Malvern, UK) instrument. Acuitas will provide the LNP used in this work to academic investigators who would like to test it. The RNA-loaded particles were characterized and subsequently stored at –80°C at an RNA concentration of ∼1 μg μl^-1^. diameter of mRNA-LNPs was ∼80 nm with a polydispersity index of 0.02-0.06 and an encapsulation efficiency of ∼95%. The DNA sequences of nucleoside-modified mRNA are listed in the Supplementary Table 1.

### Tissue sources and immunohistochemistry on frozen sections

Livers were collected directly in 4% paraformaldehyde (PFA) for overnight fixation at 4°C prior to OCT frozen block processing. For cryopreservation, tissues were washed thrice with PBS, dipped in 15% sucrose for 15 min, and then transferred to and kept in 30% sucrose solution until they sank to the bottom. The liver lobes were then cut in half lengthwise and embedded in OCT. 5 μm liver sections were cut using CM1950 Leica cryostat and slides were stored at −20°C until required for staining.

For tissues from AAV8-TBG-STAT5bCA experiments, livers were isolated into 70% ethanol for paraffin embedding. Livers were then dehydrated with 30 min of 90% ethanol then 100% ethanol, twice each, then 100% xylene. Then livers were switched to 1:1 ratio of xylene and paraffin at 60°C for 30 min, then vacuum baked in 100% paraffin for 30 min 3x at 60°C. Paraffin blocks were then sectioned with a Leica microtome, and rehydrated with 10 minutes Histoclear/xylene, then 2 x 5 minutes 100% ethanol, then 1 minute each of 90% ethanol, 70% ethanol, 50% ethanol, then water, when ready for staining.

### Liver dissociation and single-cell RNA sequencing

Liver was perfused according to the previously published method with minor modifications^41,66,83^. All solutions were pre-warmed to 40 °C and delivered at a rate of 4 ml/min. Mice were anaesthetized with ketamine/xylazine and then perfused by cannulation with a 24-gauge catheter through the inferior vena cava with 30 mL 1X Liver Perfusion Medium (Gibco by Life Technologies), while the portal vein was cut to allow the perfusate to flow out. In total, 15 mL of Earle’s Balanced Salt Solution (EBSS) containing 10mM Hepes (Ca++ and Mg++, pH 7.4) was then perfused, and finally followed with 30 mL of Liver Digest Medium (Gibco by Life Technologies) to allow complete dissociation of liver cells in situ. Livers were then extracted and mechanically dissociated in 10 mL of Liver Digest Medium. Cells were filtered through 100 μm cell strainer and filtrate was centrifuged at 50 × g for 2 min at 4 °C to obtain hepatocytes in the pellet while the supernatant was collected as the fraction of NPCs. The hepatocyte pellet was resuspended in wash media (Hepatocyte Wash Medium by Gibco + 0.1 mg/mL DNAse 1 + 10% FBS) and saved. The cell fraction caught in the 100 μm strainer was further digested in NPC Digest Medium (2.5 mg/mL Collagenase IV + 0.1 mg/mL DNAse 1), filtered through a 40 μm strainer, then centrifuged at 300 × g for 5 min at 4 °C. The supernatant was discarded, pellet was resuspended in wash media, and solution was added to the other collection of NPCs. This NPC fraction was used for the single-cell RNA sequencing analysis.

Cell concentration was counted via hemocytometer, in triplicate, then the NPC fraction was pelleted, and resuspended in a buffer of PBS + 10% FBS to a concentration of 1000 cells/uL, as per 10X Genomics cell prep protocol. Cell concentration after resuspension was checked again via hemocytometer to confirm. Single cells were captured for sequencing library preparation at the BU scRNA Seq facility using the Chromium Single Cell 3’ platform (10X Genomics). Barcoded sequencing libraries were loaded on a NextSeq500 (Illumina) with a custom sequencing setting (26 bp for Read 1, 98 bp for Read 2), to obtain a mean sequencing depth ranging from 20K to 70K mean reads per cell. All four samples were sequenced in parallel to minimize batch-to-batch variability. We excluded from analysis cell doublets, cells containing more than 25% of mitochondrial RNA reads and cells with less than 300 genes detected (indicative of dying cells). We targeted 3,000 cells per sample. However, based on the sex and treatments, the actual numbers of cells sequenced were 1688 for PBS-treated female, 756 for APAP-treated female, 605 for PBA-treated male, and 1772 for APAP-treated male. Samples were normalized using the Seurat package^84^, with scaling and correction for unwanted sources of variation, like cell degradation, as measured by the percentage of mitochondrial reads in each cell. We used PCA for linear dimensionality reduction and then used the first 20 principal components for identifying cell clusters in the sample, using the Louvain algorithm. Cell cycle scoring and other molecular signature enrichments were computed using the method from Tirosh et al.^85^. Differential expression tests were run using MAST^86^, with gene filters to reduce the burden of multiple test corrections (min.pct = 0.25, logfc.threshold = 0.25). Contrasts in factorial analyses were also run with limma/edgeR. The individual datasets were subsequently combined in Seurat and re-analyzed using the same defaults as for the individual analyses. A targeted gene set enrichment analysis was done using molecular signatures of interest from the database MsigDB (in particular from the Hallmark and Biocarta collections). Pathways of interest were scored in each cell using the same module scoring method outlined above^85^, and t-tests were used to assess differences in gene set scores between groups for each cell type. The scores were displayed in violin plots. Further exploratory analyses were performed after importing the data into SPRING^42^ for interactive visualization and further data exploratory analyses. Samples were subplotted by cell type to identify the most highly enriched genes by cell type, which were then analyzed with ENRICHR for pathway analysis, and ranked by z-score of enrichment as defined in the Bioplanet 2019 pathways database.

The raw and processed scRNA-seq fastq files are available at GEO at NCBI, with GEO accession number ##### (in process).

### ALT assay

For serum assays, blood was isolated from the inferior vena cava or tail tip, then centrifuged at 3000 x g for 15 min to separate the serum from the blood cells. Assays were performed using the Pointe Scientific kit (A7526-450) for testing serum ALT levels following manufacturers protocol. Briefly, 10 μl of serum was mixed with supplied reagent mix at 37 °C and readings were measured at 340 nm every 1 min for 5 min using Molecular Devices SpectraMax® i3x Multi-Mode microplate reader.

### Hematoxylin & eosin (H&E) assay

Histology was performed on frozen-fixed liver tissue sections. Briefly, slides were hydrated with tap water, stained with Gill’s Hematoxylin, blued with ammonia, washed in ethanol, stained with 0.25% Eosin Y, then cleared with Histoclear. Slides were mounted with permanent mounting media and observed under bright-field microscope.

### TUNEL assay

Apoptotic and necrotic cells were visualized and quantified using the Invitrogen Click-iT™ Plus TUNEL Assay for In Situ Apoptosis Detection (Alexa Fluor™ 488 dye). The protocol was adapted for use on frozen-fixed tissue sections by scaling up reagent proportions for larger volumes. Sections were counter-stained with DAPI, mounted, and observed under a fluorescence microscope green channel (Nikon Eclipse Ni-E microscope).

### Histological analyses

Images from stained sections were visualized and measured at 40X magnification on ImageJ (FIJI) using the area measurement tool. 2-3 lobes were measured and averaged per mouse.

### ELISA assay

Human GH concentration in the serum and liver tissue homogenates were measured with Human Growth Hormone Quantikine ELISA Kit (R&D Systems cat# DGH00). Serum was diluted 1:1000 in diluent provided, while tissue was diluted 1:100 in diluent provided. Liver tissue was isolated from the right median lobe and mechanically dissociated in 1mL of sterile PBS for analysis. Murine IGF1 was measured in plasma collected from a lateral tail vein nick prior to APAP injection, using the Mouse/Rat ELISA Kit (22-IG1MS-E01, ALPCO, Salem, NH).

### Statistics and reproducibility

The statistical analyses for the graphs were carried out using two-sided student’s t test for unpaired comparisons with GraphPad Prism. The statistical analyses for the Kaplan-Meier survival curves were carried out using the Log-rank Mantel-Cox test with GraphPad Prism. A p value < 0.05 was considered significant, ***<0.0001, **<0.001, *<0.05, n.s. or no result = not significant. The results are presented as the means ± standard error or standard deviation as indicated in legends.

## SUPPLEMENTAL FIGURES

**Supplemental Figure 1:**
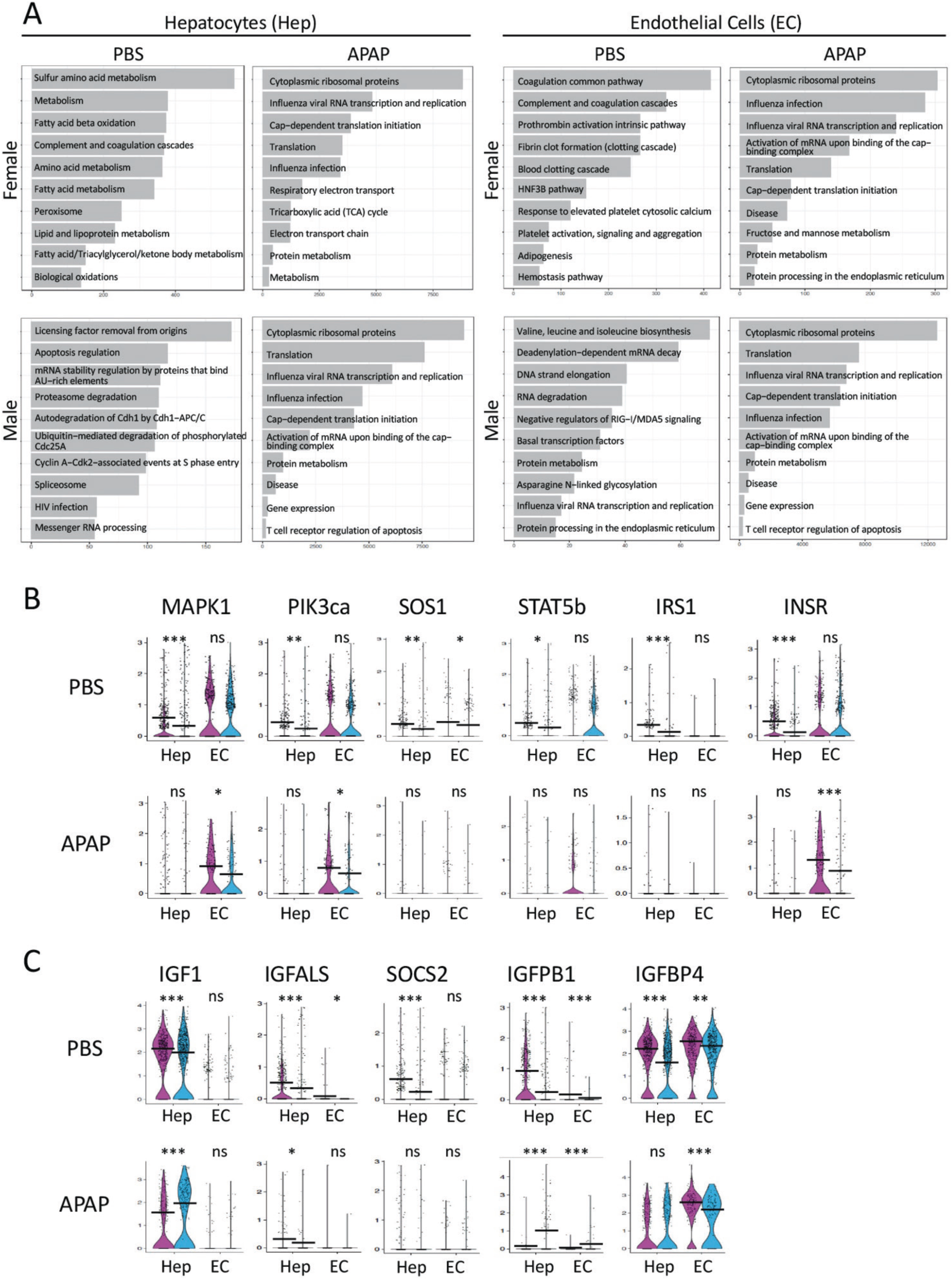
Male and female hepatocytes and endothelial cells are transcriptionally distinct. A: Single liver hepatocytes (Hep) and endothelial cells (EC) from males and females treated with PBS or APAP harvested as described in Figure 2 were analyzed for pathway enrichment using ENRICHR. From the SPRING plots, the list of the most enriched genes in pre-identified hepatocytes and endothelial cells in each group were defined, analyzed with ENRICHR for pathway analysis, and ranked by z-score of enrichment as defined in the Bioplanet 2019 pathways database. B/C: Violin plots from Heps and ECs of genes included in the BioCarta GH pathway activation (B) as well as of additional downstream genes key for GH pathway activation (C). T-tests were used to assess differences in gene set scores between groups for each cell type, p-values *<0.05, **<0.001, ***<0.0001.

**Supplemental Figure 2:**
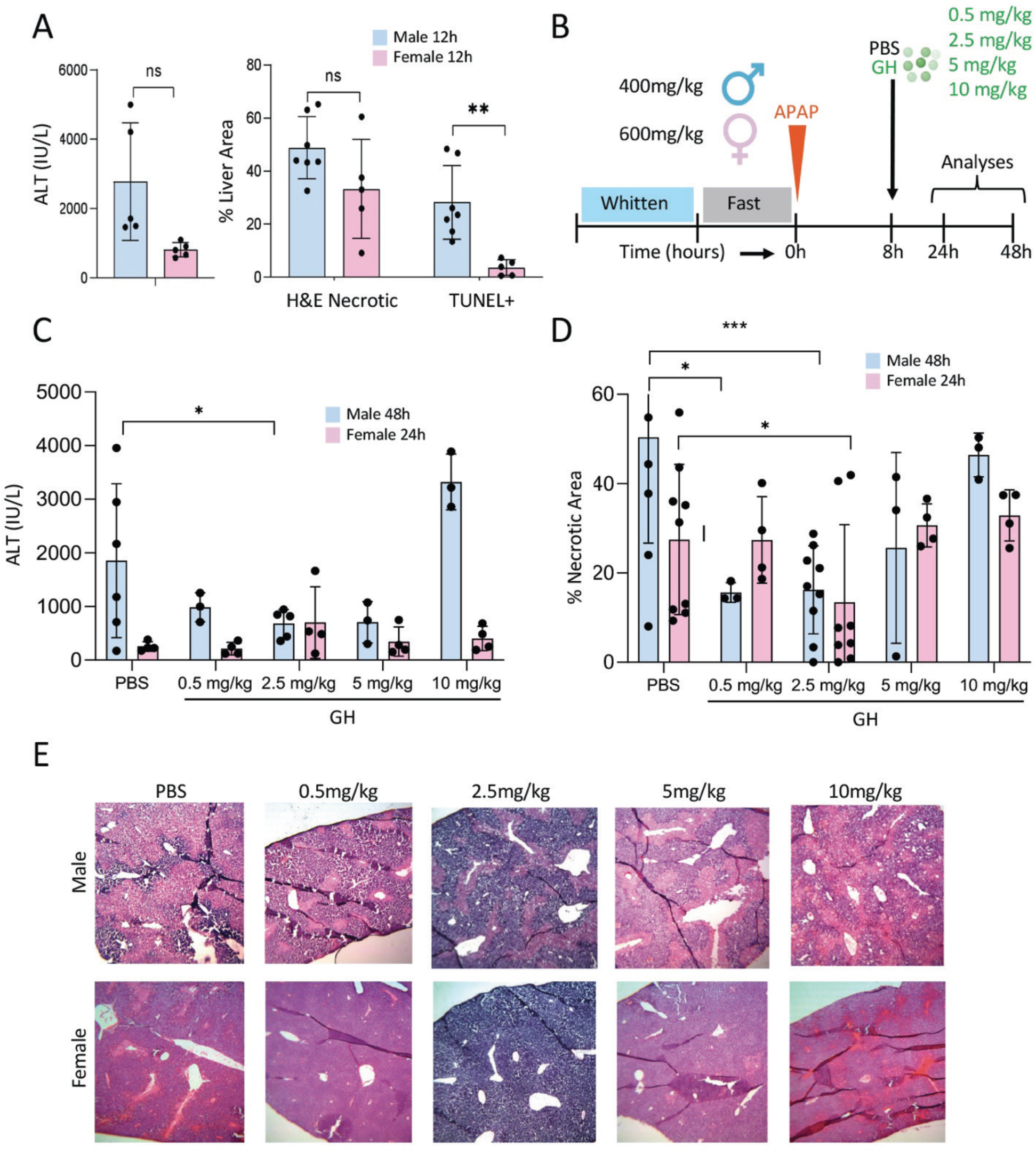
Median dosage of 2.5 mg/kg GH is most beneficial for treatment post APAP injection. A: Serum ALT, % necrotic liver tissue and % TUNEL+ liver tissue quantified from H&E stain and TUNEL stain, respectively, from males (blue) and females (pink) 12 hours post-APAP treatment with doses of 400 mg/kg and 600 mg/kg, respectively. B: Injury and treatment scheme for male and female mice injured with sex-specific doses of APAP and treated 8 hours later with 4 different doses of subcutaneous GH protein injection or PBS vehicle control, and sacrificed at either 24 hours post-treatment (females) or 48 hours post-treatment (males). C: Serum ALT levels of GH-treated and PBS-treated male and female mice 24 and 48 hours after administration of APAP. D: % Necrotic liver tissue quantified via H&E stain for males (blue) and females (pink) following PBS or GH protein treatment at 4 different doses. E: Representative H&E images for each sex at each GH dosage shown at 40x magnification. Statistics calculated via twosided student’s t test for unpaired comparisons; p-values *<0.05, **<0.001, ***<0.0001.

**Supplemental Figure 3:**
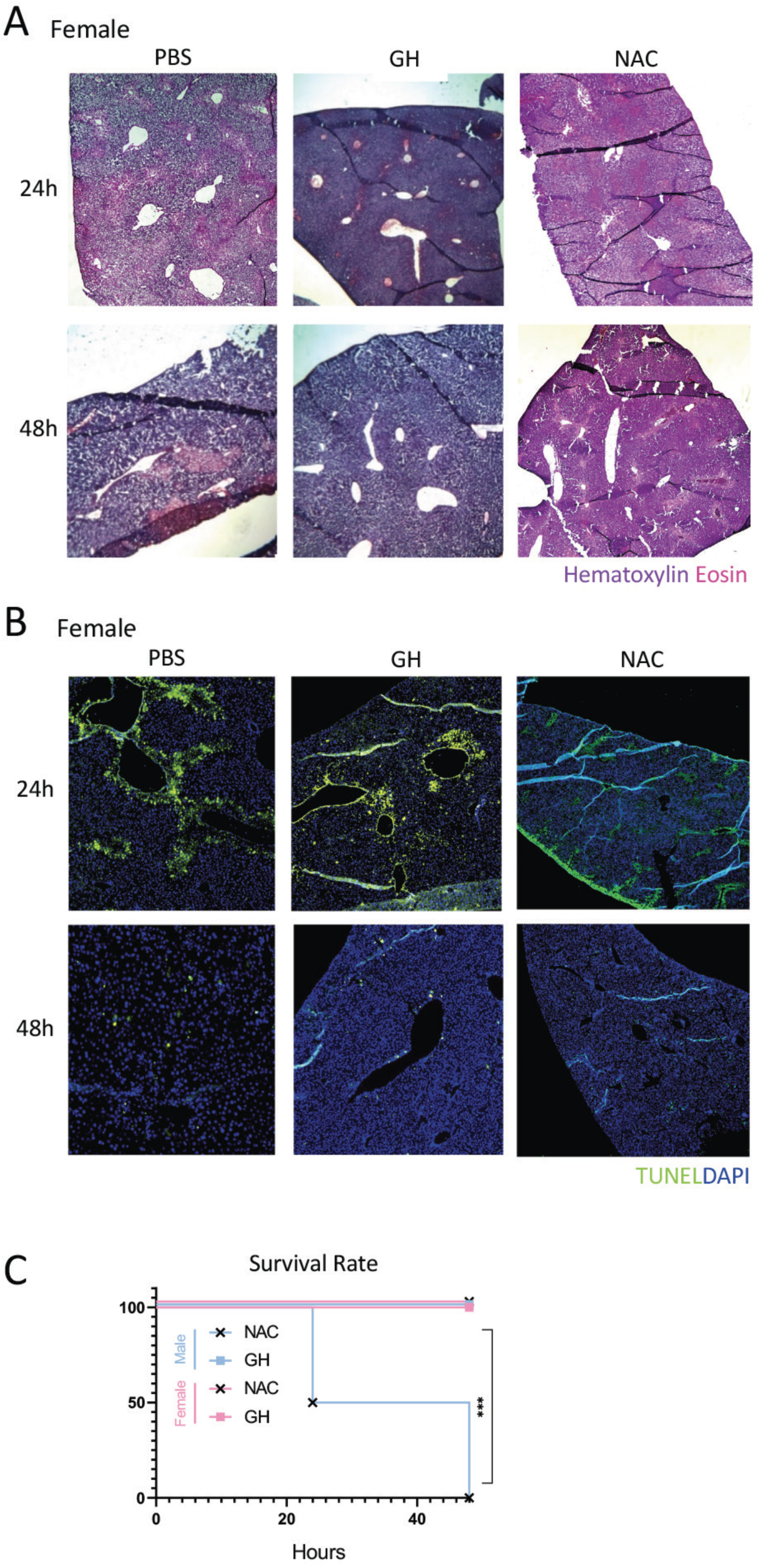
GH treatment promotes recovery to a lesser extent in females. A/B: Pictures of livers from female mice described in Figure 3 treated with APAP, NAC or PBS. Representative H&E stains (A) or TUNEL stains (B) from mice used in Figure 3 of livers per treatment, and time point post-APAP at 40X magnification. C: Survival rate calculated from GH-treated (squares; males in blue, females in pink) vs NAC-treated (black x’s) mice after APAP overdose. N= 6 mice per sex/treatment. Statistical analysis carried out using the Log-rank Mantel-Cox test.

**Supplemental Table 1:**
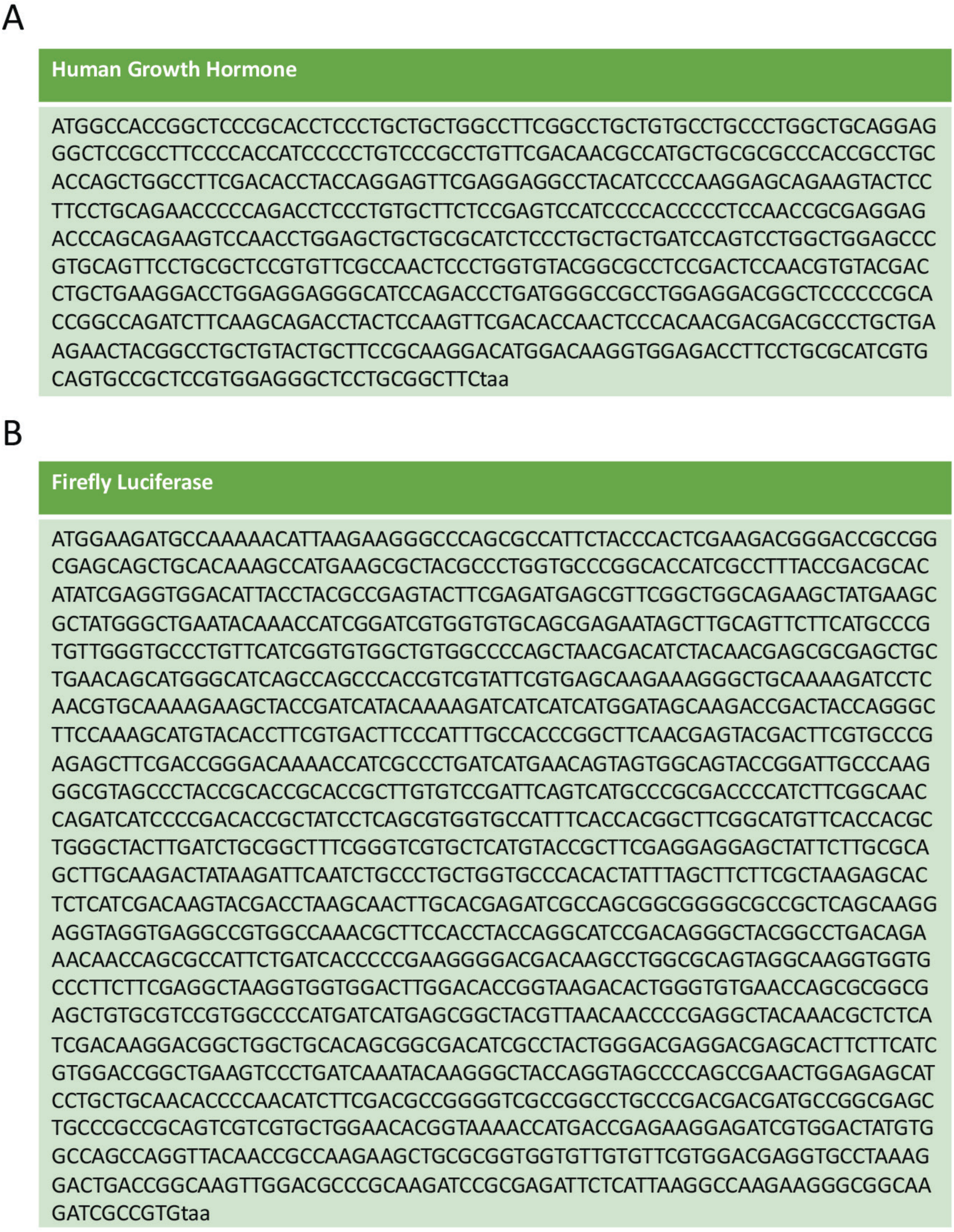
DNA sequences used as templates to in vitro transcribe nucleoside modified mRNA.

